# Mesoscopic oscillations in a single-gene circuit without delay

**DOI:** 10.1101/032029

**Authors:** N. Guisoni, D. Monteoliva, L. Diambra

## Abstract

It is well known that single-gene circuits with negative feedback loop can lead to oscillatory gene expression when they operate with time delay. In order to generate these oscillations many processes can contribute to properly timing such delay. Here we show that the time delay coming from the transitions between internal states of the *cis*-regulatory system (CRS) can drive sustained oscillations in an auto-repressive single-gene circuit operating in a small volume like a cell. We found that the cooperative binding of repressor molecules is not mandatory for a oscillatory behavior if there are enough binding sites in the CRS. These oscillations depend on an adequate balance between the CRS kinetic, and the synthesis/degradation rates of repressor molecules. This finding suggest that the multi-site CRS architecture plays a key role for oscillatory behavior of gene expression.

PACS numbers: 82.40.Bj,87.10.Mn,87.16.-b,87.16.Yc,87.18.Tt,87.18.Vf

---

Oscillatory phenomena are an essential feature of biological systems and such behavior is present at different levels of the organization of the living matter (cell, tissues, organs and individuals). At the intra-cellular level several examples of genes with oscillatory expression are known, whose periods range from ~ 40 minutes in the zebrafish somitogeneses [1] to a day in circadian clocks [2]. In general, the mechanism underlying such oscillations is a negative regulatory loop implemented in a gene-protein interaction network. The complexity of such networks vary from highly complex ones, as those described for the cell division cycle, or the circadian rhythm, to the simplest ones which were synthetically implemented in prokaryotic cells. In this sense Stricker *et al*. have shown that a synthetic single-gene circuit is able to display oscillatory behavior [3]. The dynamical system theory predicts that single-gene circuits with negative feedback loop can exhibit oscillatory gene expression when they operate with an explicit time delay [4, 5], or when such time delay is implicit in additional steps representing post-transcriptional events [6, 7]. Several processes, such as: transcript elongation, splicing, translocation, translation and phosphorylation, can contribute to generate a proper time delay. However, the individual impact of these processes on genetic clocks is not yet fully understood [8]. In this paper we show that an alternative mechanism, based on transitions between internal states of the CRS, can generate the time delay needed to induce oscillations when operates in a stochastic regime, but not in a deterministic scenario. To examine if this mechanism can generate sustainable oscillations *per se*, we devise an autorepressive single-gene loop without explicit time delay nor additional post-transcriptional events. Our model consider that gene expression is regulated by a tandem of *N* functionally identical regulatory binding sites where the gene product can bind cooperatively inhibiting its own expression (see Figure 1). This architecture has a biological counterpart in the mouse Hes1 and Hes7 genes which negatively autoregulate its own expression through three binding sites in the proximal promoter [9]. In order to emphasize the role of *cis*-regulatory system dynamics as an alternative oscillatory mechanism, we do not take into account the translation step. Also, for simplicity, we consider that the repressor synthesis occurs, at rate *a*, only when no repressor is bound to the DNA, and that they are linearly degraded at rate *g*. Thus, the deterministic reaction rate equations can be written as

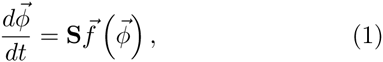

where 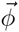 is the vector of concentrations, **S** is the stoichiometric matrix, and 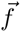 is a vector function whose component *f_j_* is the rate function of the *j*-th reaction. In our case *φ_i_* is the fraction of genes with *i* = 0,1*,…,N* bound molecules, and *Ø_n_*_+1_ is the concentration of repressors, which will be denoted hereafter by *c*. For the example of Fig. 1, when *N* = 3, there are 8 reactions, and the rate function vector is 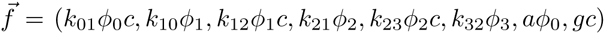, where *k_ij_* is the kinetic rate between promoter states *i* and *j*. Due to cooperativity, previous binding alters the actual binding or unbinding process. Known relationships between the system’s kinetics and thermodynamic properties allow us to write all kinetic rates, *k_s_,_s_*_+1_ and *k_s+_*_1_*,_s_* in terms of three parameters [10]: the binding rate p, the unbinding rate q, and *ɛ* which represents the cooperativity intensity, i.e., (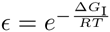, where ∆*G_I_* is the free energy of the interaction). We consider that the presence of already bound repressors alter DNA affinities increasing binding rates *k_s,s_*_+1_. Thus, following [10], we can write *k_s_,_s_*_+1_ = *ɛ^s-1^ (N* +1 – *s*) *p*, while *k_s+_*_1_*,_s_* = *s q*. An analytic exploration by standard linear analysis around the steady state of this system is only possible for *N ≤* 2. However, if the synthesis and degradation processes are much slower than the CRS kinetics, we can perform a quasi-steady state approximation on *φ_i_* variables, with *i =* 0*,…, N*, which allows to write the temporal evolution of the repressors concentration as ċ = *a F(c)* – *g c. F*(c) is a sigmoidal regulatory function with an effective half-maximum concentration *K_d_*, and effective steepness *n*. Such simplified system does not present limit cycle independently on the steepness of the monotonically decreasing regulatory function *F*.

**FIG. 1.**
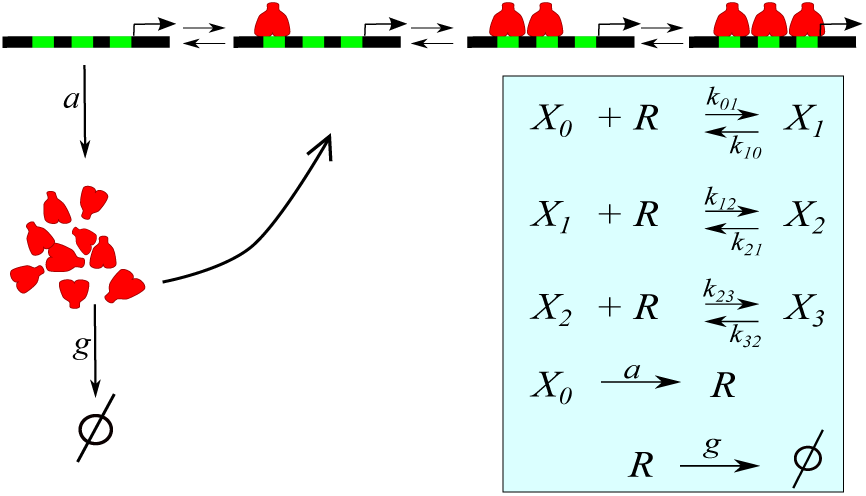
Sketch of the autorepressive single-gene loop with three binding sites. The repressor *R* (red) can bind to regulatory sites (green) on the DNA inhibiting its own synthesis. Inset: Cascade of reactions where *X_i_* represents the promoter states, and *k_i,j_* the transition rates.

A couple of simulations at different descriptive levels illustrate the importance of the role of intrinsic fluctuations on the induction of oscillations in this system. In particular, repressor concentration obtained by numerically integration of Eqs. (1) with *N* = 3, when *p* =1.7 x 10^−3^ (*μ*M min)^−1^, *q* = 0.75 min^−1^, *ɛ* = 9, *a* = 0.075 *μ*M min^−1^, and *g* = 0.3 min^−1^ ([4, 10], and references therein), presents a damped oscillation before reaching the fixed point (Figure 2A, red line), while an exact stochastic simulation of these same reactions exhibits oscillations (black line) similar to the degrade-and-fire oscillations reported in [11]. On the other hand, for lower cooperativity (Figure 2B, *ɛ* = 2), where the deterministic system does not exhibit damped oscillations, the oscillatory-like behavior of the stochastic counterpart is less apparent and more difficult to be distinguished from noise. These results show that the CRS architecture can constitutes a mechanism for noise-induced oscillations (NIO).

As we have shown, the macroscopic equations (1) are not suitable to describe correctly the phenomena occurring within a cell. In fact, inside the cells, the copy numbers of the reacting molecules are low enough for molecular discreteness to become relevant, leading to the necessity of a stochastic description of the system [12, 13]. To this end we will consider the strategy of the linear-noise approximation (LNA), developed in [14], to compute the approximate power spectrum of the number fluctuations. Within the classical LNA, the fluctuations 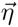 around the mean concentrations (1) are given by [15]:

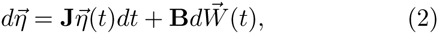

where **J** and **D** = **BB***^T^* are the Jacobian and diffusion matrices, respectively, and 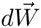 is a Wiener process. The closed-form equation for the power spectrum of fluctuations for the chemical species *i* is given by [14]:

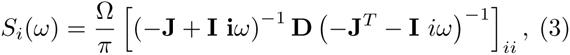

where Ω is the volume. Matrices **J** and **D** can be computed from the rate equations and from the stoichiometry matrix as 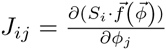, and 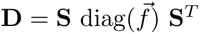 [15].

**FIG. 2.**
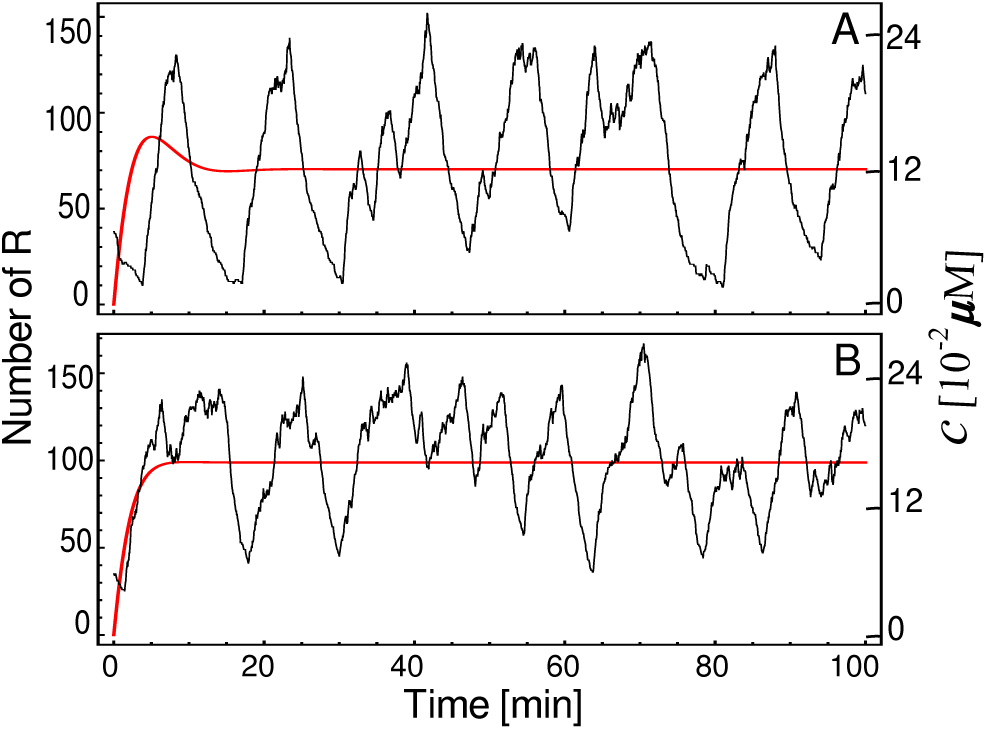
Temporal course of the concentration *c* (red lines), obtained by numerical integration of Eq. (1), and stochastic trajectories of the same system displaying an oscillatory behavior (black line). The parameter values are *N* = 3, *p* = 1.7 x 10^−6^ *μ*M min^−1^, *q* = 0.75 min^−1^, *a* = 0.075 *μM* min^−1^, and *g* = 0.3 min^−1^. In addition *ɛ* takes value of 9 for case (A) and 2 for case (B). For stochastic simulations we consider the cell volume as Ω = 1 x 10^−15^ *l*.

**FIG. 3.**
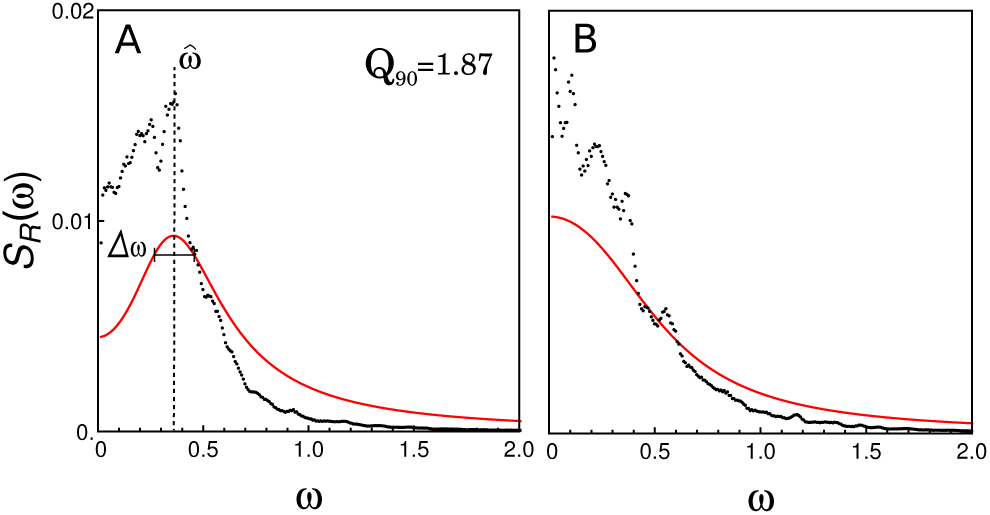
Normalized power spectral density of the fluctuations of repressor obtained from stochastic simulations of Fig. 2 (black dots), and the approximate normalized power spectral density computed by using Eq. (3) (red curve). 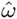 indicates the peak frequency, and ∆*ω* is the difference of the two frequencies at which the power takes the 90% of the peak value. The frequency *ω* is given in radians per minute (rad min^1^).

Figure 3 shows the normalized numerical spectrum (black dots) estimated from the exact stochastic simulations displayed in Fig. 2 and, for comparison, the corresponding power spectrum computed by Eq. (3) (red curve). The numerical spectrum was calculated by averaging the periodograms of 8000 realizations of 820 min length, and then normalizing by the total power. The numerical spectrum show peaks, which are more evident in the case of *ɛ* = 9 (Fig. 3A), where there is a stable focus. On the other hand, the approximated power spectral densities (red curve) exhibit only one peak, at *ω* = 0.37, in the case of *ɛ* = 9, in good agreement with the power spectrum computed from stochastic simulations. However, the LNA does not reflect the small peaks that also appear with *ɛ* = 2 (Fig. 3B). The presence and form of a peak in the solid curve help to distinguish sustained oscillation from apparently random fluctuations. Therefore, we will consider an oscillation quality measure [14], 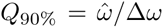, where 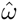 is the peak frequency of NIO, and ∆*ω* is the difference between the two frequencies at which the power takes its 90% of the peak value. In the cases of Fig. 3, the quality factor for the sustained oscillation case (Fig. 3A, *ɛ* = 9) is *Q*_90_% = 1.87, while it is not defined for the case with lower cooperativity (Fig. 3B, *ε* = 2). Increasing *p*-value 100-fold in case of *ε* = 9 also leads to oscillations. The periods of the oscillations in Fig. 3 (~ 15 min) are smaller when contrasted with the fastest oscillations in eukaryotes, however by including other time-consuming steps in the model, as elongation, protein synthesis and translocation, the oscillation period can increase significatively.

Beyond showing evidence on NIO phenomenon in a single-gene circuit without explicit delay, we want to study two key features. The first one is the steepness of the regulatory function, which is determined both by the intensity of the cooperative binding *ɛ*, and the number of regulatory binding sites *N*. The second one is the interplay between the CRS kinetic and the rate of synthesis and degradation processes. To unshroud the influence of this relationship, we scaled the rates of the synthesis and degradation processes with a parameter λ, writing *a* = λ*a*_0_, and *g* = λ*g*_0_. Thus, increasing λ speeds up the synthesis/degradation processes in relation to the CRS, but without altering the mean number of repressor molecules nor the resulting regulatory function.

Figure 4 depicts the peak frequency 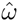 of the oscillatory behavior, and the corresponding quality factor *Q*_90_% as function of ɛ and λ for the oscillations of number fluctuation of repressor. For the parameter values studied here (the same as in Fig. 2 with *a_0_* = 0.075 *μ*M min^−1^ and *g_0_* = 0.3 min^−1^) the oscillation period ranges from 3 to 15 minutes, reaching the maximum frequency at high rates of the synthesis/degradation processes (left panel). However, in this regime the quality of oscillations are poor, as one can observe in the right panel. The *Q*_90%_-factor shows that there is a particular range for the synthesis/degradation rates, around λ = 2, where the NIO become particularly evident. Fig. 4B also shows that the auto-regulatory circuit proposed here can not present clear oscillatory behavior for slow rates of synthesis/degradation processes (λ < 2). One also can observe from Fig. 4B that increasing *ɛ* improves the NIO phenomena.

**FIG. 4.**
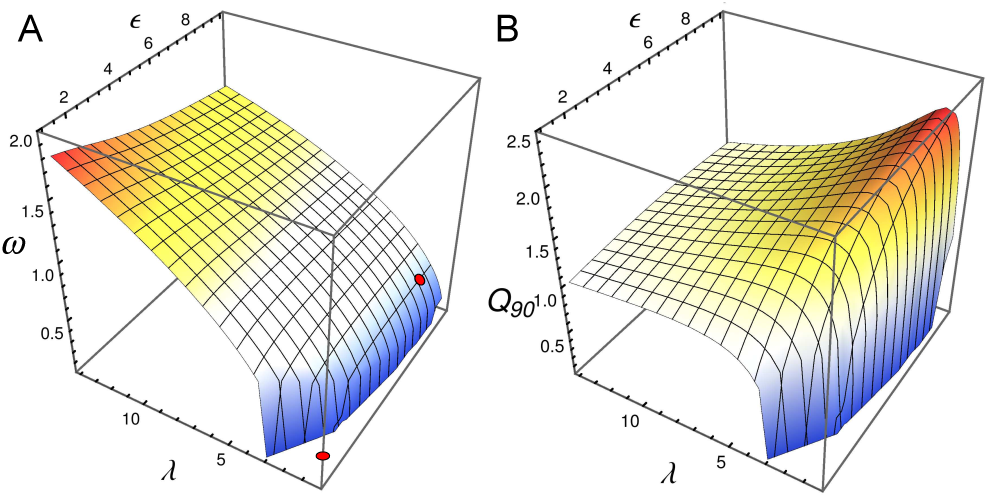
Peak frequency 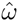 (panel A) and the quality factor *Q*_90_% (panel B) as function of *ɛ* and λ for *ɛ* for the oscillations of number fluctuation of repressor. Red dots correspond to the parameter values used in Figs. 2 and 3.

We have also studied what happens when we increase the steepness of the effective regulatory function by increasing the number of binding sites, keeping constant all other parameters. For the case of *N* = 5, one observes a slight decrease of the peak frequency when increasing *ɛ*, as shown in Fig. 5. On the other hand, as expected, the *Q*_90_%-factor increases markedly when the number of regulatory binding sites increases. In this sense, for λ = 1 and moderate *ɛ* values the *Q*_90_%-factor reaches a value of 3. An interesting feature shown in Fig. 5B is that *Q*_90%_ - factor does not increase monotonically with *ɛ*. This behavior can be linked with the fact that fluctuation levels, and probably the width of the peak frequency, increases with *ɛ* [16].

**FIG. 5.**
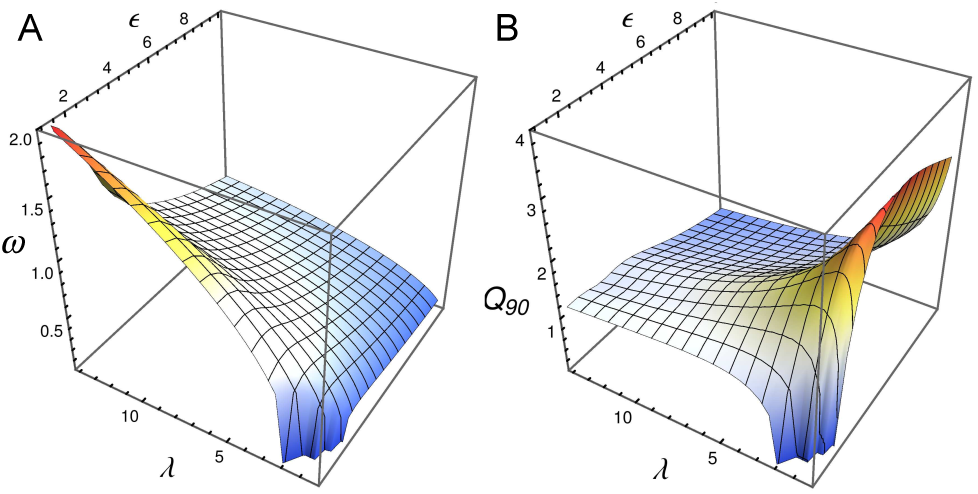
Peak frequency 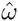 (panel A) and the quality factor *Q*_90_% (panel B) as function of *ɛ* and λ for *ɛ* for the oscillations of number fluctuation of repressor for the same parameter than Fig. 4 but with *N* = 5.

As illustrated by the previous examples, auto-repressive single gene circuit with multi-site CRS can exhibit oscillatory behavior in a mesoscopic regime without the necessity of explicit time lagged variables. The oscillatory behavior illustrated above can be explained in terms of transitions between the internal states of CRS, which are able to buffer time in the same way as additional post-transcripcional steps. In our model the time-buffering capability is determined by the mean number of sites occupied when there is a high number of repressor molecules. Consequently, for a given CRS kinetics the expression level of repressors that reach the system could be an important parameter for NIO development. To illustrate this idea we consider the same parameters for CRS as in Fig. 5, but increasing *a_0_* and decreasing *g_0_* parameters by the same factor, in order to increase the expression level. Fig. 6 shows that the peak frequency 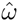 is almost the same as Fig. 5A, while the quality factor *Q*_90_% gives clear evidence that the oscillatory behavior becomes much more apparent. Remarkably, in this regime it is possible to find good oscillations even in the absence of cooperative binding (i.e., ɛ = 1).

**FIG. 6.**
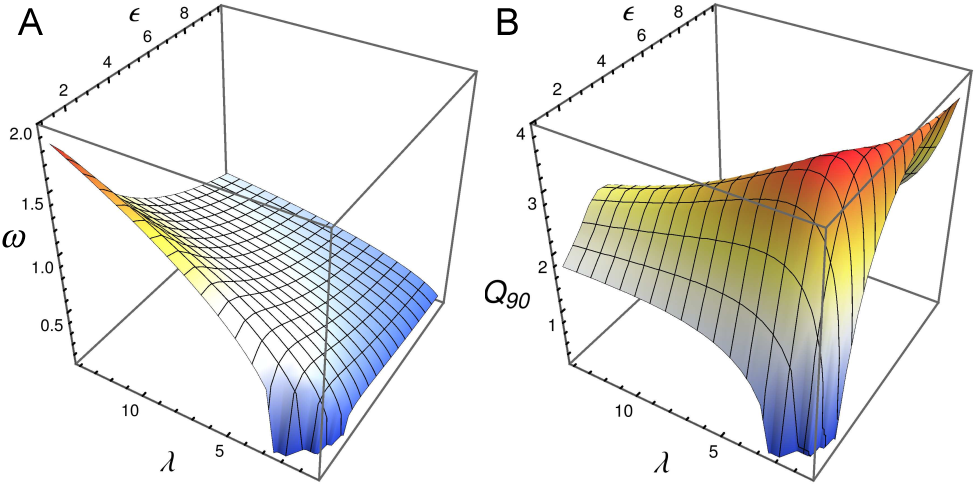
Peak frequency 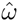 (panel A) and the quality factor *Q*_90_% (panel B) as function of *ɛ* and λ for *ɛ* for the oscillations of number fluctuation of repressors for the same parameter as Fig. 2.

In summary, we have presented a model that considers a multi-site CRS architecture and allows to unravel the role of the different elements that conform the circuit (binding sites, cooperative interactions, kinetic rates) in the development of an oscillatory behavior. Our stochastic analysis reveals that oscillations in the expression level are feasible in a broad range of parameters, while the corresponding macroscopic reaction rate equations predict the existence of stable fixed points. Strikingly, we found that the cooperative interaction, although enhances oscillations as it is already known, is not an essential ingredient because we found oscillatory behavior even in a total absence of cooperativity. On the other hand, we show that by increasing the expression level of the system, one can increase the implicit delay, derived from the multi-site CRS architecture, improving the quality of the sustained oscillations.

Our finding emphasize the role of the multi-site CRS architecture for naturally occurring genetic oscillators, such as genes Hes1 and Hes7. The expression of these genes is inhibited by their own products which can bind to three regulatory binding sites [1], as proposed here. We have shown that in a multi-site promoter the activation/deactivation process contributes to the overall time and can drive oscillation *per se*. However, though the periods of the oscillations reported here are smaller than the period of the segmentation clocks, we expect that longer periods can be reached by including in the model other time-consuming processes [8, 17, 18].

We believe that the use of detailed, and biologically interpretable, CRS in combination with stochastic analyses, offers new insights into the nature of these oscillations, especially in the context of segmentation clocks, as well as potentially aiding in the design of new synthetic biological prototypes.

We thank Alejandra Ventura and Andrés McCarthy for critical reading of the manuscript. NG and LD are members of CONICET (Argentina). DM acknowledges support from CIC (Argentina). This work was supported by CONICET, PIP #: 0020 and 0143.

